# ABO blood group is involved in the quality of the specific immune response

**DOI:** 10.1101/2021.05.23.445114

**Authors:** Sergio Gil-Manso, Iria Miguens Blanco, Bruce Motyka, Anne Halpin, Rocio Lopez-Esteban, Veronica A. Perez-Fernandez, Diego Carbonell, Luis Andrés López-Fernández, Lori West, Rafael Correa-Rocha, Marjorie Pion

## Abstract

Since December 2019, the coronavirus disease 2019 (COVID-19), caused by the severe acute respiratory syndrome coronavirus 2 (SARS-CoV-2), has spread throughout the world. To eradicate it, it is crucial to acquire a strong and long-lasting anti-SARS-CoV-2 immunity, by either natural infection or vaccination. We collected blood samples 12–305 days after positive polymerase chain reactions (PCRs) from 35 recovered individuals infected by SARS-CoV-2. Peripheral blood mononuclear cells were stimulated with SARS-CoV-2-derived peptide pools, such as the Spike (S), Nucleocapsid (N), and Membrane (M) proteins, and we quantified anti-S immunoglobulins in plasma. After 10 months post-infection, we observed a sustained SARS-CoV-2-specific CD4+ T-cell response directed against M-protein, but responses against S- or N-proteins were lost over time. Besides, we demonstrated that A-group individuals presented significantly higher frequencies of specific CD4+ T-cell responses against Pep-M than O-group individuals. The A-group subjects also needed longer to clear the virus and they lost cellular immune responses over time, compared to the O-group individuals, who showed a persistent specific immune response against SARS-CoV-2. Therefore, the S-specific immune response was lost over time, and individual factors determine the sustainability of the body’s defences, which must be considered in the future design of vaccines to achieve continuous anti-SARS-CoV-2 immunity.

**Summary:** This work describes that cellular responses against SARS-CoV-2 M-protein can be detected after 10 months but were lost against S- and N-proteins. Moreover, the individual factors; ABO-group and age influence the sustainability of the specific humoral and cellular immunity against SARS-CoV-2.

## Introduction

Since December 2019, a new severe acute respiratory syndrome coronavirus 2 (SARS-CoV-2), causing coronavirus disease 2019 (COVID-19), has spread worldwide, triggering various clinical manifestations in infected patients, such as dry cough, fatigue, fever, diarrhoea, and pneumonia(Wang et al., 2020). The SARS-CoV-2 pandemic poses a serious health threat to the global population. The most effective way to protect the population would be to achieve widespread anti-SARS-CoV-2 immunity, after either natural infection or vaccination. Strong anti-SARS-CoV-2 immunity is crucial for reducing the spread of the virus, and information about the immune system’s sustainability or efficiency in fighting the virus is central to improving patient management (Greenhalgh et al., 2020; Pascarella et al., 2020). Indeed, even people with mild symptoms may experience long-term sequelae and, possibly, immune dysregulation, and it is unknown if these long-term symptoms could be associated with re-infection or future pathogenesis (Carfì et al., 2020; Huang et al., 2021; Rubin, 2020).

Markers of the protective humoral response, such as total anti-SARS-CoV-2 immunoglobulins and neutralizing antibodies, have been observed to decrease in convalescent individuals, even though a potential long-lasting humoral B-cell memory subset was detected (Beaudoin-Bussières et al., 2020; Long et al., 2020; Ogega et al., 2020). The loss of humoral immunity has been associated with increased cases of COVID-19 recurrence (Gousseff et al., 2020; SeyedAlinaghi et al., 2020; To et al., 2020). These recurrences can be due to re-infection or viral re-activation; in both cases, immunity is at the centre of viral clearance.

Less is known about long-term cellular protection, which is pivotal for resolving viral infections and developing long-lasting immunity. Positive and promising results have suggested that cellular immunity can be generated during SARS-CoV-2 infection (Bilich et al., 2021a; Kim et al., 2020; Peluso et al., 2021), as demonstrated in SARS coronavirus infection, where memory T-cells could be detected 11 years after infection (Ng et al., 2016). The detection of these specific T-cells comprises evidence for potential pre-existing immunity mediated by T-cells cross-reactive to human common-cold coronaviruses, which might protect against SARS-CoV-2 infection (Grifoni et al., 2020; Le Bert et al., 2020; Prévost et al., 2020). Induced T-cell immunity also appears to play a critical role in SARS-CoV-2 clearance, with studies reporting strong T-cell responses in acute infection up to the convalescence phase (Grifoni et al., 2020; Sekine et al., 2020). Therefore, we studied the persistence of the antigen-specific response in individuals that had recovered from COVID-19, along with possible individual factors related to the duration and intensity of the immune response. Such information could help in the stratification of individuals according to re-infection risk factors, in order to prioritize those at high risk for immunization.

## Results

### Study participants

A total of 35 individuals were recruited following recovery from COVID-19, including 29 patients with mild symptoms, two patients with moderate symptomatology, and four asymptomatic cases, according to the WHO Working Group on the Clinical Characterisation and Management of COVID-19 infection (infection, 2020). The donors had documented dates for PCR positivity (PCR+) and/or PCR negativity (PCRneg). At the time of the study, they no longer presented symptoms related to COVID-19 (Table 1 shows the participants’ characteristics). No significant differences in gender or age were noted between the asymptomatic, mild, and moderate individuals. A total of 94.3% of the subjects were never hospitalized for COVID-19, while 5.7% were hospitalized with moderate symptoms (n = 2), none of whom required intensive care unit care (Table 1). The subjects’ ages ranged from 25 to 62 years (Table 1).

**Table 1:** Demographic and clinical characteristics of the COVID-19 convalescent patients. * From World Health Organization (WHO) Working Group on the Clinical Characterisation and Management of COVID-19 infection. (infection, 2020) †Date of SARS-CoV-2 negative PCR result was not available for nine patients.

| Characteristics | Patients (n=35) |
| --- | --- |
| **Age (years), median (range)** | 40 (25-62) |
| **Gender, n (%)** |  |
| Male | 11 (31.4) |
| Female | 24 (68.6) |
| **Blood Type, n (%)** |  |
| A (Rh+/Rh-) | 15 (42.9)/1 (2.9) |
| B (Rh+/Rh-) | 2 (5.7)/1 (2.9) |
| AB (Rh+/Rh-) | 2 (5.7)/0 (0.0) |
| O (Rh+/Rh-) | 10 (28.6)/4 (11.4) |
| **Comorbidities, n (%)** |  |
| Current smoker/ex-smoker | 3 (8.6)/4 (11.4) |
| Asthma | 3 (8.6) |
| Heart disease | 2 (5.7) |
| Obesity | 1 (2.9) |
| Hypertension | 1 (2.9) |
| Epilepsy | 1 (2.9) |
| Psoriasis | 1 (2.9) |
| Sleep apnea | 1 (2.9) |
| Fibromyalgia | 1 (2.9) |
| Diabetes | 0 (0.0) |
| Liver disease | 0 (0.0) |
| Kidney disease | 0 (0.0) |
| **WHO clinical progression scale, n (%)\*** |  |
| Mild, asymptomatic (1) | 4 (11.4) |
| Mild, symptomatic independent (2) | 29 (82.9) |
| Moderate, no oxygen therapy (4) | 1 (2.9) |
| Moderate, oxygen by mask or nasal prongs (5) | 1 (2.9) |
| **Symptoms during COVID-19, n (%)** |  |
| Fatigue | 19 (54.3) |
| Myalgia | 19 (54.3) |
| Anosmia | 16 (45.7) |
| Fever ($\geq 38$) | 14 (40.0) |
| Headache | 14 (40.0) |
| Ageusia | 13 (37.1) |
| Cough | 13 (37.1) |
| Diarrhea | 10 (28.6) |
| Dyspnea | 9 (25.7) |
| Arthralgia | 5 (14.3) |
| Nausea or vomiting | 5 (14.3) |
| Fever ($< 38$) | 3 (8.6) |
| Pneumonia | 3 (8.6) |
| Dizziness | 3 (8.6) |
| Tachycardia | 3 (8.6) |
| Sore throat | 2 (5.7) |
| Conjunctivitis | 1 (2.9) |
| Congestion | 1 (2.9) |
| Skin rash | 1 (2.9) |
| **Treatment, n (%)** |  |
| Antibiotics | 8 (22.9) |
| Hydroxychloroquine | 6 (17.1) |
| Nonsteroidal anti-inflammatory drug | 6 (17.1) |
| Anticoagulants | 4 (11.4) |
| Steroids | 2 (5.7) |
| Antiretroviral therapy | 1 (2.9) |
| **Days from PCR+ to PCRneg, median (range)†** | 17 (6-30) |
| **Days from PCR+ to data analysis, median (range)** | 154 (12-305) |

### SARS-CoV-2 specific memory T-cells in COVID-19 patients

To study the generation of SARS-CoV-2-specific memory T-cells against structural nucleocapside (N), spike (S) and membrane (M) proteins, the intracellular cytokine expression of the donor’s PBMCs was analysed after 6 hours of stimulation with peptide pools (Fig. 1A shows the gating strategy for the CD4+ subset). The studied cytokines were intracellular IL-2, IL-4, IL-17A, IFN-γ, and TNF-α, when PBMCs were non-treated (NT) or stimulated with Pep-S, Pep-M and Pep-N, cytomegalovirus (Pep-CMV), or CytoStim as a positive control for cellular activation (Fig. S1). The CMV stimulation results represented in all the figures were derived only from CMV-seropositive individuals (n = 21) screened using the IgG anti-CMV ELISA kit. The frequencies of IFN-γ- and TNF-α-producing CD4+ T-cells were significantly higher in stimulated conditions than in NT for all the peptides, demonstrating the presence of SARS-CoV-2-specific cells in almost all the individuals (Figs. 1B and 1C, respectively). However, Pep-N did not induce a significant increase in the frequency of TNF-α-producing CD4+ T-cells (Fig. 1C). As the frequency of cytokine-expressing cells was close to the NT condition in some individuals, we calculated the stimulation index (SI), for each individual, by dividing the frequency of specific T-cell response against peptides pools by the respective response in the NT control. An SI above 2 was considered to indicate a detectable response, while that below 2 corresponded to a lack of response from the individual. We observed that most individuals presented a clear and robust signal after stimulation with Pep-S, Pep-M, and Pep-CMV, but that for Pep-N was less intense (Figs. 1D and 1E). On the other hand, neither SARS-CoV-2-derived nor CMV-derived peptides induced IL-17A and IL-4 responses in CD4+ or CD8+ T-cells, even though some individuals presented an IL-2-producing CD4+ T-cell SI greater than 2 when stimulated with Pep-M (Supplemental Fig. 2A). In terms of an SI above 2 for individuals responding to the peptides, 71.4% and 65.7% of individuals responded to Pep-S, and 80% and 88.6% responded to Pep-M, while only 62.9% and 45.7% showed a detectable response to Pep-N, according to CD4+ T-cells (considering IFN-γ and TNF-α, respectively; see Fig. 1F). Strikingly, fewer than 50% of individuals presented responses to any peptide pools derived from SARS-CoV-2 in the CD8+ T-cells (Fig. 1G). We assume that this result was not derived from experimental bias for CD8+ T-cells, as good responses were observed for the CMV-derived peptide pool and CytoStim (Supplemental Fig. 2B).

**Figure 1:**
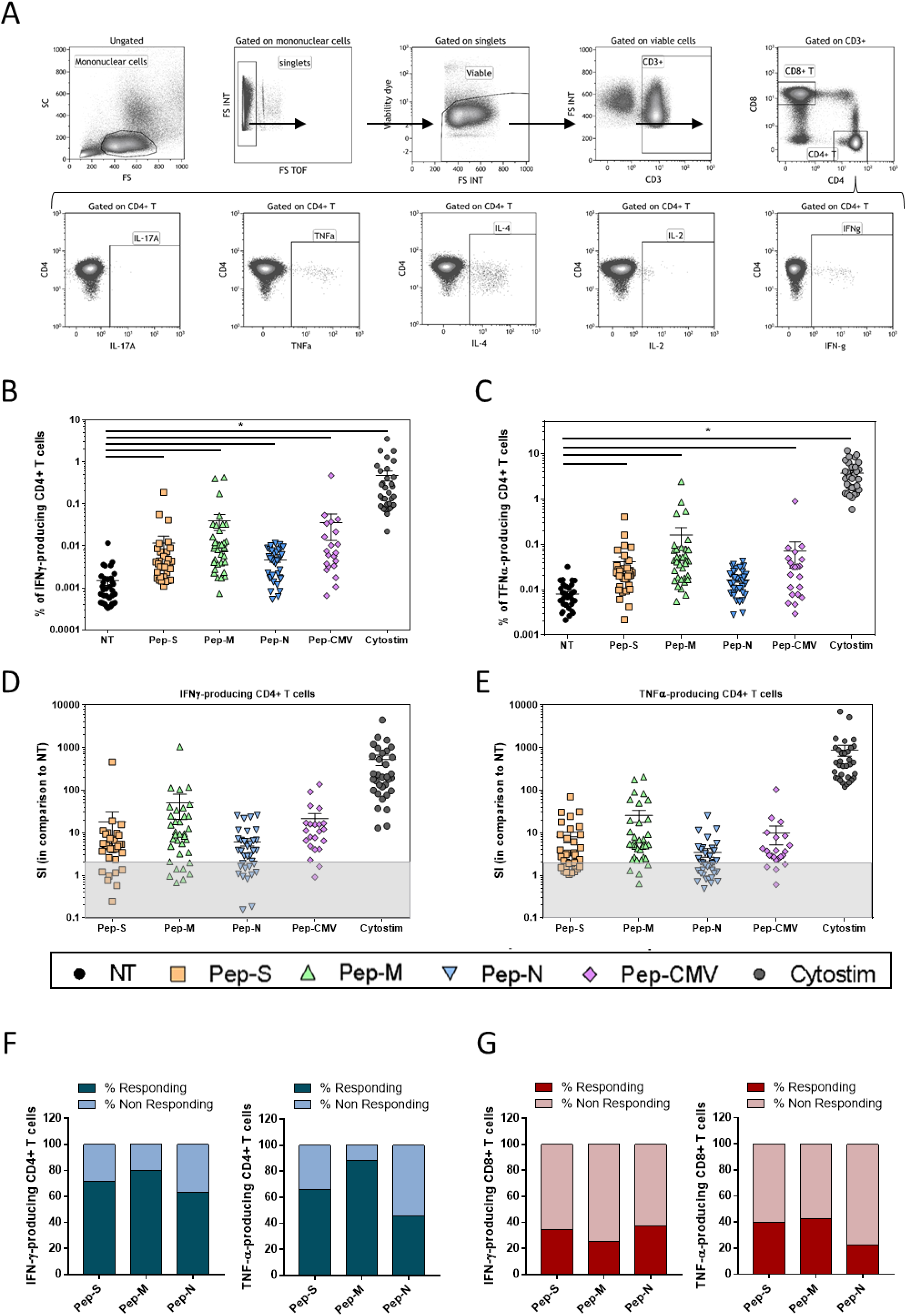
Gating strategy and specific memory T-cell responses to SARS-CoV-2-derived peptide pools in 35 convalescent patients. (A) Gating strategies to define SARS-CoV-2-specific CD4+ T-cells. Representative examples of flow cytometry plots of SARS-CoV-2-specific CD4+ T-cells after 6 h stimulation with Spike (Pep-S). PBMCs were isolated and stimulated by SARS-CoV-2-derived peptide pools (Pep-S, Pep-M, and Pep-N), with CMV-derived peptides (Pep-CMV), or with CytoStim. IL-17A, TNF-α, IL-4, IL-2, and IFN-γ expression was detected intracellularly, by flow cytometry. (B) Frequencies of IFN-γ-producing CD4+ T-cells. (C) Frequencies of TNF-α-producing CD4+ T-cells. (D) Stimulation index (SI) of IFN-γ-producing CD4+ T-cells. (E) SI of TNF-α-producing CD4+ T-cells. The SI for each subject was calculated by dividing the frequency of cytokine-producing CD4+ T-cells for the stimulated condition (Peptides or CytoStim) by the frequency of cytokine-producing CD4+ T-cells in the non-treated condition (NT). Each symbol corresponds to an individual. The grey area represents an SI lower than 2, which is considered to indicate a negative response to the stimulation, in comparison to NT. One-way ANOVA with multiple-comparison Kruskal–Wallis tests were used. *p < 0.05. (F) Stacked bars comparing the frequency of individuals with a specific T-cell response when cells were stimulated by Pep-S, Pep-M, or Pep-N. Individuals with SIs greater than 2 were considered responding individuals, and those with SIs lower than 2, as non-responding individuals. The response was observed as the intracellular IFN-γ or TNF-α production in CD4+ cells. (G) Stacked bars comparing the frequencies of individuals responding and not responding to the SARS-CoV-2-derived peptides in CD8+ T-cells.

**Figure 2:**
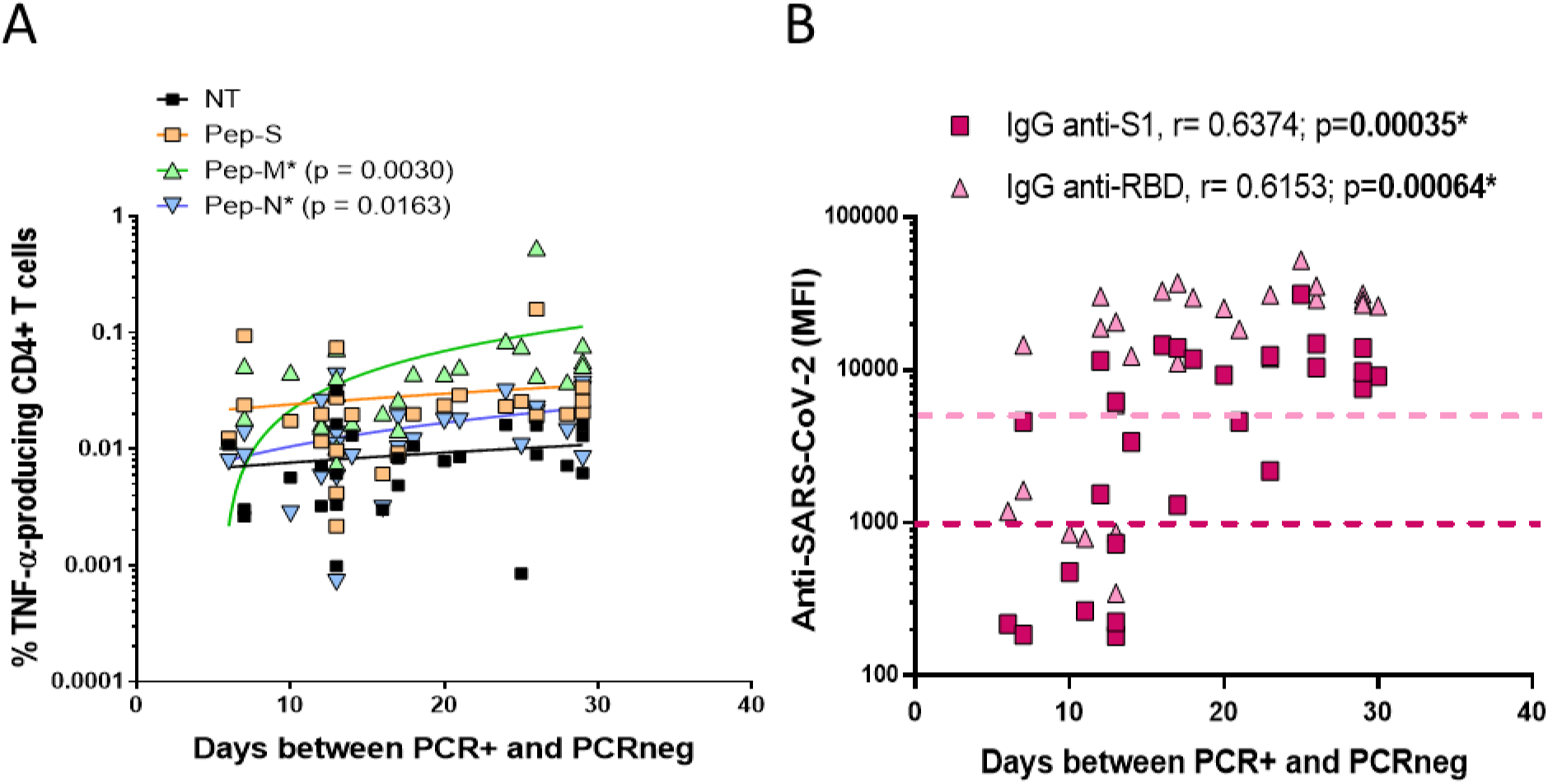
Frequencies of TNF-α-producing cells and time necessary for viral clearance. Correlation between frequencies of (A) TNF-α-producing CD4+ T-cells when non-stimulated (NT) or stimulated with Pep-S, Pep-M, or Pep-N, and the time between PCR+ and PCRneg. PCR+ corresponds to the detection of the SARS-CoV-2 infection, while PCRneg corresponds to the first PCR negative after the infection. Therefore, both dates allow the time for viral clearance to be determined. (B) Correlation between plasma levels of IgG anti-S1 and anti-RBD (anti-SARS-CoV-2 immunoglobulins) and the time between PCR+ and PCRneg. Coloured dotted lines estimated threshold of positivity for anti-SARS-CoV-2 immunoglobulin detection. Correlations were assessed using Spearman’s rank correlation; *p < 0.05 was considered significant. Each symbol corresponds to an individual.

In summary, Pep-M induced the strongest CD4+ T-cell memory responses, followed by Pep-S and, finally, Pep-N. Moreover, SARS-CoV-2-derived peptides induced responses in almost all the individuals tested, but mostly of the CD4+ memory T-cell type.

### Detection of SARS-CoV-2-specific memory T-cells, according to the time of viral clearance and time post-infection

Our study included individuals with histories of COVID-19, who were recruited 12 to 305 days after testing positive for SARS-CoV-2 by PCR+ (P-PCR+; up to 10 months). Two of the individuals analysed did not respond to any of the SARS-CoV-2-derived peptides. These two individuals were asymptomatic at the moment of PCR detection, with low viral load showed by high cycle thresholds (CTs) in real-time PCR.

First, we correlated the time between the beginning of the infection (the first PCR+ after the appearance of symptoms) and the end of the infection (the first PCRneg after COVID-19), corresponding to the period needed for viral clearance, with the frequency of cells responding to SARS-CoV-2-derived peptide pools. Patients recovered from the viral infections 6–30 days after the detection of the virus. The longer the time of active infection, the higher the TNF-α-specific response to Pep-M and Pep-N (p = 0.0030 and p = 0.0163, respectively; Fig. 2A). It has already been noted that anti-SARS-CoV-2 immunoglobulins decrease in convalescent individuals after several months (Bilich et al., 2021b); however, it is unknown whether the time of viral clearance is also crucial for the generation of anti-SARS-CoV-2 antibodies. The longer the time of active infection, the higher the levels of plasma IgG anti-S1 and IgG anti-RBD immunoglobulins (p = 0.00035 and p = 0.00064, respectively; Fig. 2B).

We then followed the evolution of anti-SARS-CoV-2 antibodies and specific T-cell responses over time post-infection. Even if IgG anti-S1 and anti-RBD were still detectable in the plasma after 10 months post-infection, their levels diminished over time. Specifically, the decrease in IgG anti-RBD was significant (p = 0.00614; Fig. 3A). On the other hand, the SI values of the TNF-α-producing CD4+ T-cells for Pep-S and Pep-N were negatively correlated over time (p = 0.0184 and p = 0.0194, respectively; Fig. 3B). We then divided the individuals into two groups: one regrouping individuals 12–150 days P-PCR+ (recent infection, up to 5 months post-infection) and one regrouping individuals 150–305 days P-PCR+ (late infection). In the recent infection group, 81.2%, 87.5%, and 68.7% of the individuals responded to Pep-S, Pep-M, and Pep-N, respectively, in terms of the frequency of TNF-α-producing CD4+ T-cells. However, the individuals in the late infection group presented 47.4%, 89.5%, and 26.3% rates of response to the same peptide pools. Therefore, the frequency of individuals with TNF-α-CD4+ specific memory T-cells against S- or N-derived peptides diminished as time passed after infection (Fig. 3C). However, it was encouraging to observe that the frequency of individuals with TNF-α-CD4+ memory against the M-protein remained identical, regardless of the time P-PCR+.

**Figure 3:**
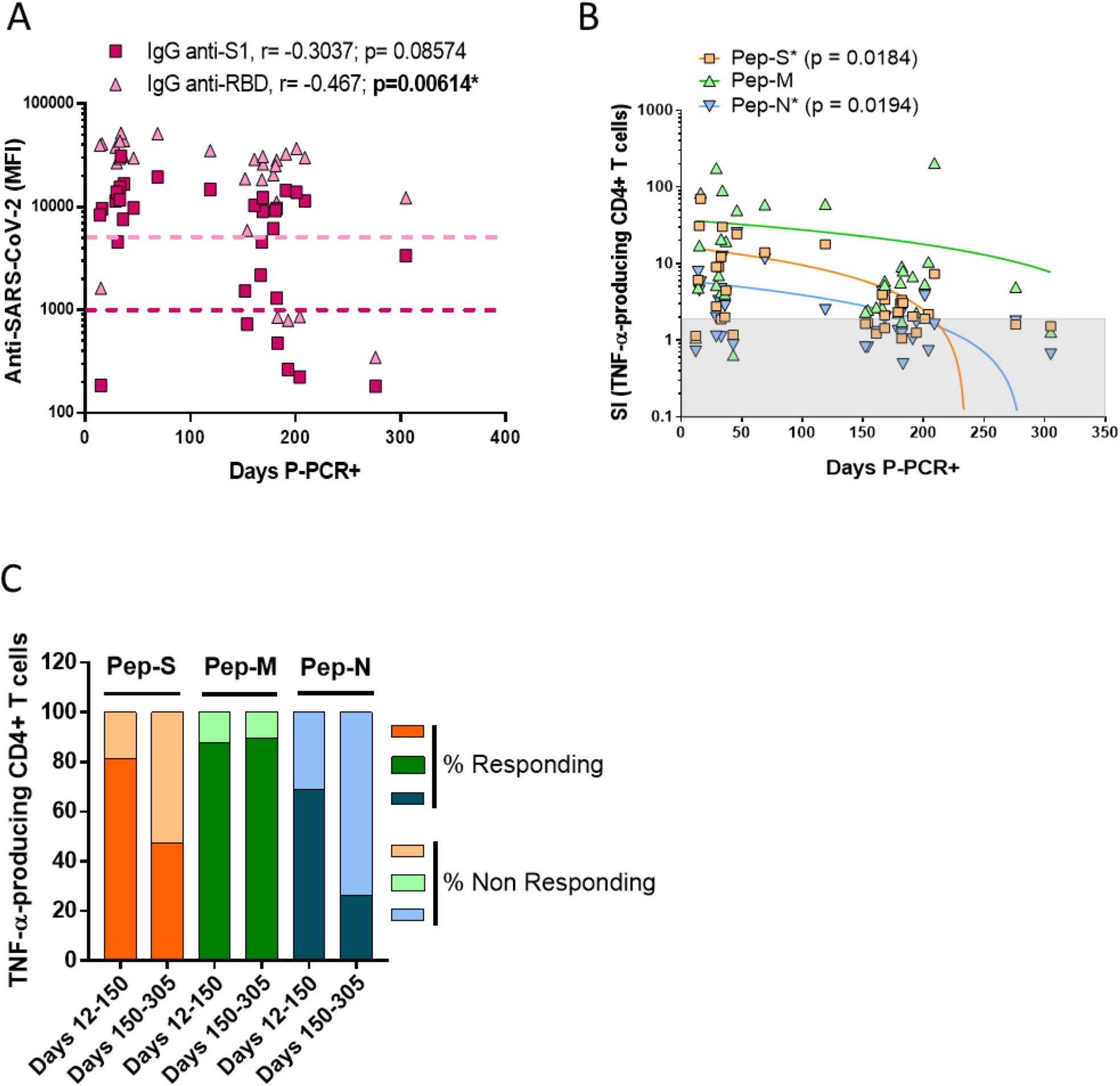
Specific CD4+ T-cell response to SARS-CoV-2 peptides at 10 months post-infection. (A) Correlation between the plasma levels of anti-S1 and anti-RBD (anti-SARS-CoV-2 immunoglobulins) and the time between the detection of COVID-19 infection (PCR+) and the time of sample processing (days post-PCR+; P-PCR+). Coloured dotted lines represent the threshold of detection. (B) Correlation between the SI of TNF-α-producing CD4+ T-cells after stimulation and days P-PCR+. Symbols in the grey zones represent samples with SIs less than 2, indicating non-responding individuals. Correlations were assessed using Spearman’s rank correlation; * p < 0.05 was considered significant. Each symbol corresponds to an individual. (C) Stacked bars represent the frequencies of individuals with TNF-α-specific T-cell responses corresponding to SIs ≥ 2 (responding) or < 2 (non-responding) when cells were stimulated by Pep-S (orange), Pep-M (green), or Pep-N (blue) in individuals tested 12–150 days P-PCR+ or 150–305 days P-PCR+.

### Individual factors associated with viral clearance and severity of symptoms

In our study, most of the individuals did not have comorbidities, and almost all had mild symptoms (Table 1). We analysed whether some other factors associated with infection susceptibility, such as age and ABO group, were related to a better response to SARS-CoV-2. A positive correlation was observed between age and the time needed to reach viral clearance (Fig. 4A; p = 0.0280). In our study, 40% of the individuals belonged to group O (n = 14), 45.5% to group A (n = 16), 8.5% to group B (n = 3), and 6% to group AB (n = 2). These frequencies were in agreement with those found in the general Spanish population. Due to the low number of individuals having the B and AB blood groups, we focused our analysis on A-group versus O-group. The A-group individuals needed a median of 23 days to reach viral clearance, while the O-group individuals needed a median of 13 days (p = 0.0229; Fig. 4B).

**Figure 4:**
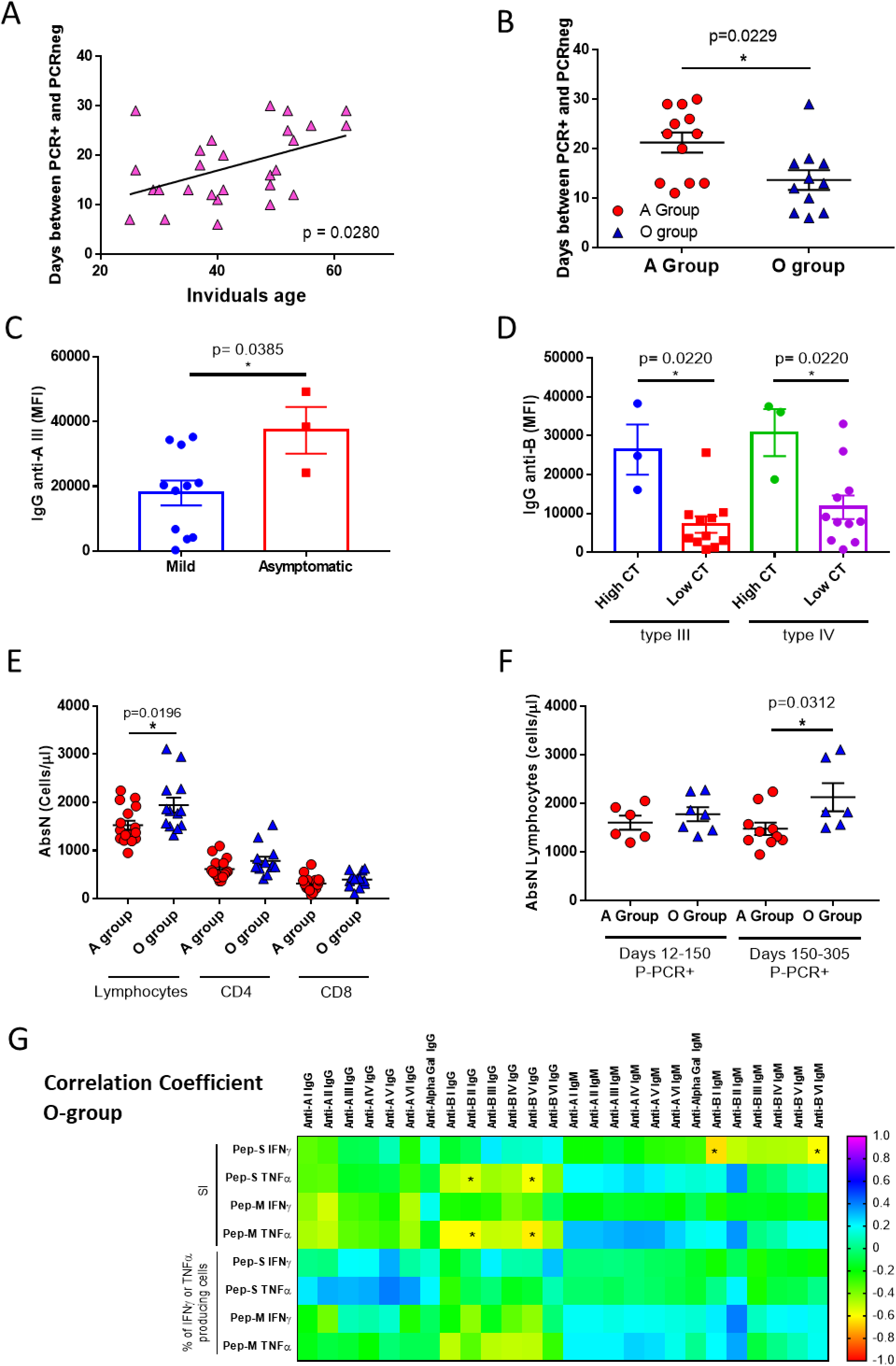
Age and blood groups as factors for viral clearance and absolute numbers of lymphocytes. (A) Correlation between the ages of the individuals and numbers of days P-PCR+. Correlations were assessed using Spearman’s rank correlation. Each symbol corresponds to an individual. (B) Numbers of days between PCR+ and PCRneg in A-group and O-group individuals. Mann– Whitney U test. (C) Plasma levels of anti-A type III immunoglobulins in mild and asymptomatic COVID-19 individuals. Mann–Whitney U test. (D) Plasma levels of anti-B type III and IV immunoglobulins in individuals presenting high and low CT in the real-time PCR the day of sample processing in mild and asymptomatic COVID-19 individuals. Mann–Whitney U tests. (E) Absolute numbers of lymphocytes, CD4+ T-cells, and CD8+ T-cells in A- and O-group individuals. Mann–Whitney U tests. (F) Absolute numbers of lymphocytes in A- and O-group individuals tested 12–150 days P-PCR+ or 150–305 days P-PCR+. Mann–Whitney U tests. (G) Heat map of Spearman correlation coefficients for indicated features in O-group individuals. *p < 0.05 was considered significant.

Then, we analysed whether the level of anti-A or anti-B immunoglobulins found in the O-group individuals could be related to the severity of COVID-19 symptoms. Of the four asymptomatic patients recruited, three belonged to the O-group, and, even though the number of individuals was low, we observed that the asymptomatic individuals showed significantly higher levels of IgG anti-A (type III) in the plasma than those with mild symptoms (Fig. 4C). Additionally, the levels of IgG anti-B (type III and IV) were significantly higher in the O-group individuals with low viral loads, on the day of sample processing (as detected by real-time PCR, high CT), than those with higher viral loads (low CT; Fig. 4D).

It has previously been observed that the severity of symptoms and viral load are associated with immune dysregulation. We observed that O-group individuals showed higher absolute numbers (AbsN) of total lymphocytes (1862 ± 174 cells/µl, mean ± SEM) than those in the A-group (1440 ± 93 cells/µl, mean ± SEM; p = 0.0196; Fig. 4E). This can be explained by the lymphopenia already observed in COVID-19 individuals, even those with mild symptoms. Additionally, the individuals from the O-group seemed to recover the AbsN of total lymphocytes after more than 150 days P-PCR+ (1783 ± 143 cells/µl in recent infection group versus 2132 ± 290 cells/µl in late infection group, mean ± SEM; p = 0.0312). Individuals from the A-group presented even lower AbsN of lymphocytes after more than 150 days P-PCR+ (1607 ± 145 cells/µl in recent infection group versus 1487 ± 142 cells/µl in late infection group, mean ± SEM; Fig. 4F). Therefore, ABO grouping might influence the course of infection. Indeed, negative correlations were observed between the levels of anti-B immunoglobulins and the level of IFN-γ- and TNF-α-producing CD4+ T-cell responses, when activated with Pep-S or Pep-M, in the O group (SI or frequencies of specific CD4+ T-cells; Fig. 4G), indicating that, the lower the anti-B immunoglobulins in O-group individuals, the higher the generation of specific immune responses.

### ABO group and anti-SARS-CoV-2-specific immune response

As a low AbsN of lymphocytes can cause dysregulation in the immune response, we studied whether the blood group could influence the generation of anti-SARS-CoV-2 memory in the long term post-infection. When cells were stimulated with Pep-M, the frequencies of IFN-γ- and TNF-α-producing CD4+ T-cells were significantly higher in group A than in group O (p = 0.0085 and p = 0.0245, respectively; Supplementary Fig. 3). In addition, when cells were stimulated with Pep-S or Pep-M, the SI of TNF-α-producing CD4+ T-cells significantly decreased as time passed (p = 0.0287 and p = 0.0415, respectively) in the A-group individuals, but not in the O-group (Fig. 5A). Moreover, the SI of TNF-α-producing CD4+ T-cells was also lower in the O-group than in the A-group (recent infection groups) when cells were stimulated with Pep-M (p = 0.0426; Fig. 5B). The frequencies of TNF-α-producing CD4+ T-cells were also lower in the later infection A-group than in the recent infection A-group individuals (p = 0.0075; Fig. 5B). Similar results were observed when correlating the days P-PCR+ and level of anti-RBD in the plasma in the A-group individuals (p < 0.0001; Fig. 5C). The cellular and humoral specific responses decreased as time passed from infection in the A group but not in the O group, showing that a strong but labile anti-SARS-CoV-2 immune response could occur in the A group.

**Figure 5:**
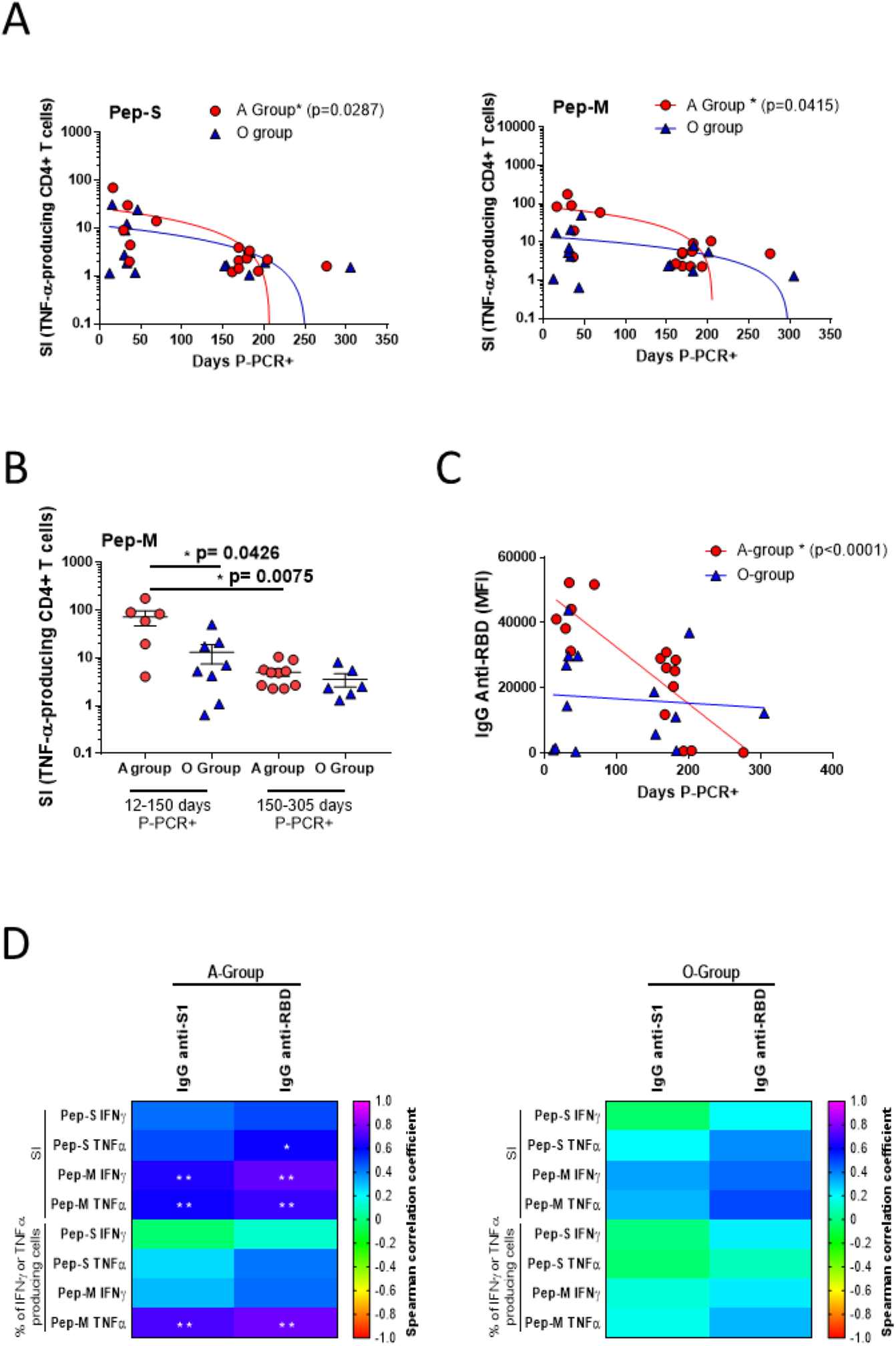
Blood groups as factor for specific CD4+ T-cell response. (A) Correlation between SI of TNF-α-producing CD4+ T-cells when stimulated with Pep-S and Pep-M in O-/A-groups and days P-PCR+. Correlations were assessed using Spearman’s rank correlation. (B) Frequencies of TNF-α-producing CD4+ T-cells, when stimulated with Pep-M in O-/A-group individuals, tested 12–150 days P-PCR+ or 150–305 days P-PCR+. Mann–Whitney U tests. (C) Correlation between level of anti-RBD immunoglobulins and days P-PCR+. Correlations were assessed using Spearman’s rank correlation. Each symbol corresponds to an individual. (D) Heat map of Spearman correlation coefficients of indicated features in A-group (left panel) and O-group (right panel) individuals. *p < 0.05; **p < 0.01.

Finally, we checked whether there were correlations between the cellular and humoral specific responses. Positive correlations between the levels of anti-SARS-CoV-2 immunoglobulins and specific cellular responses were observed in A-group individuals but not in O-group individuals, likely indicating a co-ordinated cellular and humoral immune response in the A group (Fig. 5D). In summary, the ABO blood group is an essential factor that can influence the time of viral clearance and the intensity of the TNF-α-associated response over time P-PCR+, with the A group showing the highest TNF-α-associated response, as well as a significant decrease in response intensity 10 months post-infection.

## Discussion

In this study, we aimed to evaluate the induction and duration of T-cell specific memory and humoral immunity in individuals who had recovered from SARS-CoV-2 infections with asymptomatic/mild symptoms individuals who represent the great majority of infected subjects. We also studied some susceptibility factors that may be associated with the development of sustained immunity. At 10 months post-infection, almost all the enrolled individuals were still positive for SARS-CoV-2-specific IFN-γ- and TNF-α-producing CD4+ T-cells against the M viral protein and for anti-S1 or anti-RBD immunoglobulins. However, we also observed a decreased frequency of individuals with SARS-CoV-2-specific responses against the S or N viral proteins with the time passed since the infection, as had already been observed in other studies (Breton et al., 2021; Dan et al., 2021). In addition to a decreased immunity developed against the SARS-CoV-2 S-protein, we observed that the CD8+ T-cell memory response seemed to be deficient. Whether a robust CD8+ T response might be generated is a worthwhile question, as the CD8+ cytotoxic T-cell response is generally profoundly implicated in viral clearance. It has been shown that COVID-19 subjects present elevated Th2-cytokines (IL-4 or IL-10), which could inhibit the Th1 response (Gutiérrez-Bautista et al., 2020). It has already been described that Th1 participates in the activation, proliferation, and differentiation of the cytotoxic memory CD8+ T-cells (Sercan et al., 2010). Low CD8+ T-cell activation due to an inadequate Th1 response and high Th2 response could explain the low CD8+ T-cell response frequency. Therefore, it would be interesting to design vaccines against non-spike proteins along with the S-protein, which could promote the generation of stronger CD4+ and CD8+ T-cell memory.

Looking for factors that could influence the frequency of response against SARS-CoV-2-derived peptides, we observed that the age of the subjects was important for rapid viral clearance. Older individuals are more susceptible to developing severe COVID-19 symptoms, due to several factors such as a decline in organ function, basal inflammation, or the presence of comorbidities (Mayoral et al., 2020; Zhou et al., 2020). Nevertheless, we report that age is also associated with viral clearance for the first time in this paper, to the best of our knowledge.

ABO blood groups have been implicated in the susceptibility to and severity of SARS-CoV-2 infections (Anastassopoulou et al., 2020; Padhi et al., 2020). In particular, the A group has been associated with an increased risk of acquiring COVID-19, compared to non-A groups (Wu et al., 2020; Zhao et al., 2020). In this study, we showed that blood groups influence the generation of memory T-cell responses for the first time. We showed that A-group subjects needed more time to clear the virus than the O group, and that they presented higher frequencies of IFN-γ and TNF-α CD4+ T-cell responses against Pep-M than O-group individuals. Moreover, there were positive correlations between the humoral and cellular specific responses only in A-group individuals but not in O-group individuals. These results might indicate that the A group can develop a more robust response against Pep-M; we hypothesize that this could be due to the A-group individuals needing more time to clear the virus. Thus, the longer the time of viral exposure, the stronger the response, even in mild COVID-19 cases. In severe COVID-19 cases, patients achieve a higher adaptive immune response, where a more elevated and/or extended viral load could be related to the higher immune response (Demaret et al., 2020). On the other hand, A-group individuals showed significantly fewer TNF-α-related T-cell memory responses and lower plasma levels of anti-Spike immunoglobulins after a long time P-PCR+ than the O-group subjects. Thus, even though the memory T-cell response was initially higher in the A group, the TNF-α response was lost over time, as indicated by the plasma humoral anti-SARS-CoV-2 immunoglobulins, showing a deterioration of the sustainability of the specific immune response.

One characteristic of COVID-19 is lymphopenia, even in individuals with mild symptoms, with more profound lymphopenia in patients with severe symptoms (Zhou et al., 2020). We observed that the A group presented significantly lower absolute numbers of total lymphocytes than the O group. These results might indicate that the immune systems in the A-group subjects were more affected by the infection than the O group’s immune systems. According to the literature, the O group seems to be associated with a lower risk of acquiring COVID-19 than non-O groups (Zhao et al., 2020). The principal factor relating the ABO group to COVID-19 susceptibility may be the presence of anti-A, anti-B, or anti-glycan antibodies such as anti-Gal or anti-N-Glycolyl neuraminic acid (Galili, 2020). The presence of such antibodies especially in O-group individuals could inhibit the SARS-CoV-2 S protein’s adhesion to ACE2-expressing cell lines, as has already been observed for SARS-CoV (Guillon et al., 2008). Therefore, our hypothesis is that such virus blocking may lower the infectious viral load in O-group subjects; then, one can assume that the lower viral load associated with the potential protective effect of anti-blood-group antibodies in an O-group subject accelerates the clearance of the virus and reduces its impact on the immune system. However, the implications of other factors, such as yet-unknown factors related or unrelated to ABO blood groups that could potentially affect the susceptibility to SARS-CoV-2 infection, cannot be discarded.

Taken together, these results confirmed the existence of SARS-CoV-2-specific CD4+ T-cell and humoral responses in the majority of the individuals who had recovered from COVID-19 at 10 months post-infection. However, the response generated by the virus was identified as a predominantly CD4+ T-cell over CD8+ T-cell response, with more robust responses against M- or S-peptide pools over Pep-N. A more in-depth analysis demonstrated that the intensity of the humoral and memory T-cell response is related to the ABO blood group and age. Therefore, determining the individual characteristics that may influence the immune response to SARS-CoV-2 must be considered for the future design of vaccines with long-term efficacy.

## Materials and Methods

### Patients and blood samples

Blood samples and questionnaire data regarding donor characteristics during COVID-19 infection from SARS-CoV-2 convalescent donors were collected at the General University Hospital Gregorio Marañón, Spain, from 6/2020 to 12/2020. Informed consent was obtained under the Declaration of Helsinki protocol. The study was approved by the local ethics committee and performed according to their guidelines (COV1-20-007). SARS-CoV-2 infection was confirmed by a PCR test after a nasopharyngeal swab. SARS-CoV-2 donors were recruited among health workers of the General University Hospital Gregorio Marañón in Madrid who had been infected by SARS-CoV-2 between March and December 2020. Samples were collected at a single time point, between 12 days post-positive PCR (P-PCR+) and 305 days P-PCR+ (Table 1). Whole blood was labelled for surface markers. PBMCs were isolated by density gradient centrifugation. The serum was separated by centrifugation, and the supernatant was stored at -80 °C. The classification of symptoms was based on responses to a questionnaire by individual donors. The score (Asymptomatic/Mild/Moderate) was based on the criteria of the WHO Working Group on the Clinical Characterization and Management of COVID-19 infection (infection, 2020). The characteristics of the SARS-CoV-2-recovered donors are detailed in Table 1.

### Stimulation with SARS-CoV-2 peptide pools

SARS-CoV-2 PepTivator peptide pools (Miltenyi Biotec, Bergisch Gladbach, Germany), mainly consisting of 15-mer sequences with 11 amino acids (aa), were used. The peptide pool for the Spike-protein (Pep-S) contained the sequence domains aa 304–338, 421–475, 492–519, 683– 707, 741–770, 785–802, and 885–1273. The peptide pools for the membrane glycoprotein (Pep-M) or the nucleocapside phosphoprotein (Pep-N) mainly consisted of 15-mer sequences with 11 aa overlap, covering the complete sequence of the M or N protein. Two positive controls for activation were used: PepTivator against cytomegalovirus (Pep-CMV, Miltenyi Biotec), which consisted of 15-mer peptides with 11 amino acids, covering the complete sequence of the pp65 protein of human cytomegalovirus, and CytoStim (Miltenyi Biotec), an antibody-based component that acts similarly to a superantigen of the T-cell receptor. Negative controls were left non-treated (NT). PBMCs were prepared from EDTA collection tubes (Vacutainer® K2E, BD) by gradient centrifugation. A total of 1.5 × 10^6^ PBMCs were stimulated with 1 µg/mL of peptide pools for 6 h in TexMACS™ GMP Medium (Miltenyi Biotec), supplemented with 5% AB human serum (Sigma-Aldrich, St. Gallen, Switzerland). Brefeldin A (10 µg/ml, Sigma-Aldrich) was added at the beginning of the stimulation.

### Staining for intracellular cytokines and cell surface markers

Whole blood was labelled for surface markers with the antibodies listed in Supplemental Table 1. After surface labelling, red blood cells were lysed using RBC Lysis/Fixation Solution (BioLegend, San Diego, CA, U.S.). Surface markers were analysed by flow cytometry, using a MACSQuant Analyzer 16 cytometer (Miltenyi Biotec). Peptide-specific T-cells were characterized after 6 h of stimulation by cell surface and intracellular cytokine staining. Briefly, cells were surface-stained, stained with viability dye, fixed/permeabilized, and intracellularly stained (antibodies listed in Supplemental Table 1). The cells were then analysed by flow cytometry, using a Gallios cytometer (Beckman Coulter, Nyon, Switzerland). All the cytometry data were analysed using the Kaluza software (Beckman Coulter). The gating strategy applied for the analyses of flow cytometry-acquired data is provided in Fig. 1 and Supplemental Fig. 1.

### CMV IgG Detection

The 96-well CMV IgG ELISA (Abcam, Cambridge, MA, US) was performed according to the manufacturer’s instructions. This ELISA detects human anti-cytomegalovirus IgG. The final interpretation of positivity was determined by a ratio above a threshold value given by the manufacturer: positive (ratio > 11), negative (ratio < 9), or non-defined (ratio 9–11). Quality control was performed, following the manufacturer’s instructions, on the day of testing.

### Detection of blood group antigens

To detect the presence or absence of A, B, and/or RhD antigens on red blood cells, DiaClon Anti-A, DiaClon Anti-B, DiaClon Anti-AB, and DiaClon Anti-D (Bio-Rad, Basel, Switzerland) were used, with whole blood diluted in isotonic saline solution (Braun, Hessen, Germany), according to the manufacturer’s instructions. The mix was centrifuged and then resuspended, in order to observe macroscopic agglutination.

### Testing for ABO and SARS-CoV-2 antibodies using Luminex single-antigen beads

ABO-A and ABO-B subtype glycans I–VI were conjugated to bovine serum albumin (BSA), as previously described (Jeyakanthan et al., 2016), and an optimized protein-coupling procedure was used to link each subtype antigen to individual Luminex beads (Angeloni et al., 2018). BSA-only-coupled beads were used to determine the background reactivity, while Galα1-3Galβ1-(3)4GlcNAc-R (α-Gal) BSA-coupled beads were used as a positive control (Halpin et al., 2018). Coupling was confirmed using different monoclonal antibodies, including those specific to A subtypes I–VI (clone A98, Novaclone, Immucor, Dartmouth, NS, Canada), B subtypes I–VI (clones B84 and B97, Novaclone), A/B subtype II (JTL-4) (Jeyakanthan et al., 2015), and A/B subtypes III/IV (JTL-2) (Jeyakanthan et al., 2015). Bound IgM monoclonal antibody was detected with PE-labelled goat anti-mouse IgM secondary antibody (Southern Biotech, Birmingham, AL, US) (Halpin et al., 2018).

SARS-CoV-2 S1 (Abcam) and RBD (Sino Biological, Wayne, PA, US) proteins were conjugated to Luminex beads using standard coupling procedures (Angeloni et al., 2018). Coupling was confirmed using a rabbit IgG anti-SARS-CoV-2 Spike monoclonal antibody (Sino Biological) and PE-conjugated goat anti-rabbit IgG secondary antibody (Southern Biotech).

To detect serum ABO and SARS-CoV-2 antibodies, sera (25-fold dilution) were incubated with Luminex beads for 30 minutes at room temperature, washed, and then incubated with a 50-fold dilution of PE-conjugated goat anti-human IgM or IgG (both from Thermo Fisher, Waltham, MA, US) for 30 minutes at room temperature. Samples were acquired using a FLEXMAP 3D® Luminex system (Toronto, Canada).

### Statistical analysis

Data are displayed as means with standard error. The statistical tests used to evaluate the experiments are described within the respective figure legends. Continuous data were tested for normality of distribution, and individual groups were tested by use of the Mann–Whitney U test. Spearman’s rho (r) was calculated to assess the correlation between continuous data. Graphs were plotted using the GraphPad Prism 7.00 software. Statistical analyses were conducted using GraphPad Prism 7.00 and the SPSS (IBM, version 25, Armonk, NY, US) software.

## Authorship Contributions

M.P. and R.C.R. designed research. M.P., S.G.M., R.L.E, V.A.P.F, A.H., D.C. and B.M. performed research. A.H., B.M., L.A.L.F. and L.J.W contributed vital new reagents or analytical tools. S.G.M, D.C., L.A.L.F. and I.M.B. collected data. All the authors interpreted and discussed the data. M.P. and R.C.R. wrote the manuscript. L.A.L.F., B.M., L.J.W., I.M.B, and S.G-M. revised the manuscript. All the authors read and approved the final manuscript.

## Acknowledgements

This paper is in memoriam to Prof. Alberto Tejedor and all those who succumbed while working on the global pandemic’s front line to relieve the suffering of their patients. The authors thank all the health workers who participated in this study. We acknowledge Dr. Maribel Clemente from the Cell Culture Unit and Dr. Laura Díaz from the Cytometry Unit of IiSGM. We acknowledge José Maria Bellon from the Statistical unit of IiSGM. This work was partially supported by grants from the Instituto de Salud Carlos III (ISCIII) (PI18/00506; COV20/00063), co-funded by ERDF (FEDER) Funds from the European Commission, “A way of making Europe”. This work was partially financed by the Madrid Community grant B2017/BMD3727 and the IiSGM Intramural grant PI-MP-2018. This work was partially funded by a grant from “Fundación Familia Alonso” (FFA-FIBHGM-2019). S.G-M. was supported by the Youth Employment Program, co-financed by the Madrid community and FEDER Founds (PEJ-2020-AI/BMD-17954), and by the ACT4COVID consortium. The Canadian teams were generously supported by the Stollery Children’s Hospital Foundation, through the Women and Children’s Health Research Institute, Edmonton, Alberta, Canada. The funders had no role in study design, data collection and analysis, decision to publish, or manuscript preparation.

## REFERENCES

Anastassopoulou, C., Z. Gkizarioti, G.P. Patrinos, and A. Tsakris. 2020. Human genetic factors associated with susceptibility to SARS-CoV-2 infection and COVID-19 disease severity. Hum Genomics 14:40.

Angeloni, S., S. Das, S. Dunbar, V. Stone, and S. Swift. 2018. XMAP Cookbook: A collection of methods and protocols for developing multiplex assays with xMAP technology. In Luminex Corporation, Austin, TX. 1–150.

Beaudoin-Bussières, G., A. Laumaea, S.P. Anand, J. Prévost, R. Gasser, G. Goyette, H. Medjahed, J. Perreault, T. Tremblay, A. Lewin, L. Gokool, C. Morrisseau, P. Bégin, C. Tremblay, V. Martel-Laferrière, D.E. Kaufmann, J. Richard, R. Bazin, and A. Finzi. 2020. Decline of Humoral Responses against SARS-CoV-2 Spike in Convalescent Individuals. mBio 11:

Bilich, T., A. Nelde, J.S. Heitmann, Y. Maringer, M. Roerden, J. Bauer, J. Rieth, M. Wacker, A. Peter, S. Horber, D. Rachfalski, M. Marklin, S. Stevanovic, H.-G. Rammensee, H.R. Salih, and J.S. Walz. 2021a. Differential kinetics of T cell and antibody responses delineate dominant T cell epitopes in long-term immunity after COVID-19. Cell

Bilich, T., A. Nelde, J.S. Heitmann, Y. Maringer, M. Roerden, J. Bauer, J. Rieth, M. Wacker, A. Peter, S. Hörber, D. Rachfalski, M. Märklin, S. Stevanović, H.G. Rammensee, H.R. Salih, and J.S. Walz. 2021b. T cell and antibody kinetics delineate SARS-CoV-2 peptides mediating long-term immune responses in COVID-19 convalescent individuals. Sci Transl Med 13:

Breton, G., P. Mendoza, T. Hägglöf, T.Y. Oliveira, D. Schaefer-Babajew, C. Gaebler, M. Turroja, A. Hurley, M. Caskey, and M.C. Nussenzweig. 2021. Persistent cellular immunity to SARS-CoV-2 infection. J Exp Med 218:

Carfì, A., R. Bernabei, F. Landi, and G.A.C.-P.-A.C.S. Group. 2020. Persistent Symptoms in Patients After Acute COVID-19. JAMA 324:603–605.

Dan, J.M., J. Mateus, Y. Kato, K.M. Hastie, E.D. Yu, C.E. Faliti, A. Grifoni, S.I. Ramirez, S. Haupt, A. Frazier, C. Nakao, V. Rayaprolu, S.A. Rawlings, B. Peters, F. Krammer, V. Simon, E.O. Saphire, D.M. Smith, D. Weiskopf, A. Sette, and S. Crotty. 2021. Immunological memory to SARS-CoV-2 assessed for up to 8 months after infection. Science

Demaret, J., G. Lefèvre, F. Vuotto, J. Trauet, A. Duhamel, J. Labreuche, P. Varlet, A. Dendooven, S. Stabler, B. Gachet, J. Bauer, B. Prevost, L. Bocket, E.K. Alidjinou, M. Lambert, C. Yelnik, B. Meresse, L. Dubuquoy, D. Launay, S. Dubucquoi, D. Montaigne, E. Woitrain, F. Maggiotto, M. Bou Saleh, I. Top, V. Elsermans, E. Jeanpierre, A. Dupont, S. Susen, T. Brousseau, J. Poissy, K. Faure, M. Labalette, and L.C.R.N. (LICORNE). 2020. Severe SARS-CoV-2 patients develop a higher specific T-cell response. Clin Transl Immunology 9:e1217.

Galili, U. 2020. Host Synthesized Carbohydrate Antigens on Viral Glycoproteins as “Achilles’ Heel” of Viruses Contributing to Anti-Viral Immune Protection. Int J Mol Sci 21:

Gousseff, M., P. Penot, L. Gallay, D. Batisse, N. Benech, K. Bouiller, R. Collarino, A. Conrad, D. Slama, C. Joseph, A. Lemaignen, F.X. Lescure, B. Levy, M. Mahevas, B. Pozzetto, N. Vignier, B. Wyplosz, D. Salmon, F. Goehringer, E. Botelho-Nevers, and i.b.o.t.C.s. group. 2020. Clinical recurrences of COVID-19 symptoms after recovery: Viral relapse, reinfection or inflammatory rebound? J Infect 81:816–846.

Greenhalgh, T., M. Knight, C. A’Court, M. Buxton, and L. Husain. 2020. Management of post-acute covid-19 in primary care. BMJ 370:m3026.

Grifoni, A., D. Weiskopf, S.I. Ramirez, J. Mateus, J.M. Dan, C.R. Moderbacher, S.A. Rawlings, A. Sutherland, L. Premkumar, R.S. Jadi, D. Marrama, A.M. de Silva, A. Frazier, A.F. Carlin, J.A. Greenbaum, B. Peters, F. Krammer, D.M. Smith, S. Crotty, and A. Sette. 2020. Targets of T Cell Responses to SARS-CoV-2 Coronavirus in Humans with COVID-19 Disease and Unexposed Individuals. Cell 181:1489-1501.e1415.

Guillon, P., M. Clément, V. Sébille, J.G. Rivain, C.F. Chou, N. Ruvoën-Clouet, and J. Le Pendu. 2008. Inhibition of the interaction between the SARS-CoV spike protein and its cellular receptor by anti-histo-blood group antibodies. Glycobiology 18:1085–1093.

Gutiérrez-Bautista, J.F., A. Rodriguez-Nicolas, A. Rosales-Castillo, P. Jiménez, F. Garrido, P. Anderson, F. Ruiz-Cabello, and M. López-Ruz. 2020. Negative Clinical Evolution in COVID-19 Patients Is Frequently Accompanied With an Increased Proportion of Undifferentiated Th Cells and a Strong Underrepresentation of the Th1 Subset. Front Immunol 11:596553.

Halpin, A., J. Pearcey, T.L. Lowary, C.W. Cairo, M. Jeyakanthan, B. Motyka, S. Maier, and L.J. West. 2018. Modernizing ABO antibody detection: time for new methodologies to support ABO-incompatible transplantation? ASHI Quaterly

Huang, Y., M.D. Pinto, J.L. Borelli, M.A. Mehrabadi, H. Abrihim, N. Dutt, N. Lambert, E.L. Nurmi, Chakraborty, A.M. Rahmani, and C.A. Downs. 2021. COVID Symptoms, Symptom Clusters, and Predictors for Becoming a Long-Hauler: Looking for Clarity in the Haze of the Pandemic. medRxiv

infection, W.W.G.o.t.C.C.a.M.o.C.-. 2020. A minimal common outcome measure set for COVID-19 clinical research. Lancet Infect Dis 20:e192–e197.

Jeyakanthan, M., P.J. Meloncelli, L. Zou, T.L. Lowary, I. Larsen, S. Maier, K. Tao, J. Rusch, R. Chinnock, N. Shaw, M. Burch, K. Beddows, L. Addonizio, W. Zuckerman, E. Pahl, J. Rutledge, K.R. Kanter, C.W. Cairo, J.M. Buriak, D. Ross, I. Rebeyka, and L.J. West. 2016. ABH-Glycan Microarray Characterizes ABO Subtype Antibodies: Fine Specificity of Immune Tolerance After ABO-Incompatible Transplantation. Am J Transplant 16:1548–1558.

Jeyakanthan, M., K. Tao, L. Zou, P.J. Meloncelli, T.L. Lowary, K. Suzuki, D. Boland, I. Larsen, M. Burch, N. Shaw, K. Beddows, L. Addonizio, W. Zuckerman, B. Afzali, D.H. Kim, M. Mengel, A.M. Shapiro, and L.J. West. 2015. Chemical Basis for Qualitative and Quantitative Differences Between ABO Blood Groups and Subgroups: Implications for Organ Transplantation. Am J Transplant 15:2602–2615.

Kim, D.S., S. Rowland-Jones, and E. Gea-Mallorquí. 2020. Will SARS-CoV-2 Infection Elicit Long-Lasting Protective or Sterilising Immunity? Implications for Vaccine Strategies (2020). Front Immunol 11:571481.

Le Bert, N., A.T. Tan, K. Kunasegaran, C.Y.L. Tham, M. Hafezi, A. Chia, M.H.Y. Chng, M. Lin, N. Tan, M. Linster, W.N. Chia, M.I. Chen, L.F. Wang, E.E. Ooi, S. Kalimuddin, P.A. Tambyah, J.G. Low, Y.J. Tan, and A. Bertoletti. 2020. SARS-CoV-2-specific T cell immunity in cases of COVID-19 and SARS, and uninfected controls. Nature 584:457–462.

Long, Q.X., B.Z. Liu, H.J. Deng, G.C. Wu, K. Deng, Y.K. Chen, P. Liao, J.F. Qiu, Y. Lin, X.F. Cai, D.Q. Wang, Y. Hu, J.H. Ren, N. Tang, Y.Y. Xu, L.H. Yu, Z. Mo, F. Gong, X.L. Zhang, W.G. Tian, L. Hu, X.X. Zhang, J.L. Xiang, H.X. Du, H.W. Liu, C.H. Lang, X.H. Luo, S.B. Wu, X.P. Cui, Z. Zhou, M.M. Zhu, J. Wang, C.J. Xue, X.F. Li, L. Wang, Z.J. Li, K. Wang, C.C. Niu, Q.J. Yang, X.J. Tang, Y. Zhang, X.M. Liu, J.J. Li, D.C. Zhang, F. Zhang, P. Liu, J. Yuan, Q. Li, J.L. Hu, J. Chen, and A.L. Huang. 2020. Antibody responses to SARS-CoV-2 in patients with COVID-19. Nat Med 26:845–848.

Mayoral, E.P., M.T. Hernández-Huerta, L. Pérez-Campos Mayoral, C.A. Matias-Cervantes, G. Mayoral-Andrade, L.L. Barrios, and E. Pérez-Campos. 2020. Factors related to asymptomatic or severe COVID-19 infection. Med Hypotheses 144:110296.

Ng, O.W., A. Chia, A.T. Tan, R.S. Jadi, H.N. Leong, A. Bertoletti, and Y.J. Tan. 2016. Memory T cell responses targeting the SARS coronavirus persist up to 11 years post-infection. Vaccine 34:2008–2014.

Ogega, C.O., N.E. Skinner, P.W. Blair, H.S. Park, K. Littlefield, A. Ganesan, P. Ladiwala, A.A. Antar, S.C. Ray, M.J. Betenbaugh, A. Pekosz, S. Klein, Y.C. Manabe, A.L. Cox, and J.R. Bailey. 2020. Durable SARS-CoV-2 B cell immunity after mild or severe disease. medRxiv

Padhi, S., S. Suvankar, D. Dash, V.K. Panda, A. Pati, J. Panigrahi, and A.K. Panda. 2020. ABO blood group system is associated with COVID-19 mortality: An epidemiological investigation in the Indian population. Transfus Clin Biol 27:253–258.

Pascarella, G., A. Strumia, C. Piliego, F. Bruno, R. Del Buono, F. Costa, S. Scarlata, and F.E. Agrò. 2020. COVID-19 diagnosis and management: a comprehensive review. J Intern Med 288:192–206.

Peluso, M.J., A.N. Deitchman, L. Torres, N.S. Iyer, C.C. Nixon, S.E. Munter, J. Donatelli, C. Thanh, Takahashi, J. Hakim, K. Turcios, O. Janson, R. Hoh, V. Tai, Y. Hernandez, E. Fehrman, M.A. Spinelli, M. Gandhi, L. Trinh, T. Wrin, C.J. Petropoulos, F.T. Aweeka, I. Rodriguez-Barraquer, J.D. Kelly, J.N. Martin, S.G. Deeks, B. Greenhouse, R.L. Rutishauser, and T.J. Henrich. 2021. Long-Term SARS-CoV-2-Specific Immune and Inflammatory Responses Across a Clinically Diverse Cohort of Individuals Recovering from COVID-19. medRxiv

Prévost, J., R. Gasser, G. Beaudoin-Bussières, J. Richard, R. Duerr, A. Laumaea, S.P. Anand, G. Goyette, M. Benlarbi, S. Ding, H. Medjahed, A. Lewin, J. Perreault, T. Tremblay, G. Gendron-Lepage, N. Gauthier, M. Carrier, D. Marcoux, A. Piché, M. Lavoie, A. Benoit, V. Loungnarath, G. Brochu, E. Haddad, H.D. Stacey, M.S. Miller, M. Desforges, P.J. Talbot, G.T.G. Maule, M. Côté, C. Therrien, B. Serhir, R. Bazin, M. Roger, and A. Finzi. 2020. Cross-Sectional Evaluation of Humoral Responses against SARS-CoV-2 Spike. Cell Rep Med 1:100126.

Rubin, R. 2020. As Their Numbers Grow, COVID-19 “Long Haulers” Stump Experts. JAMA

Sekine, T., A. Perez-Potti, O. Rivera-Ballesteros, K. Strålin, J.B. Gorin, A. Olsson, S. Llewellyn-Lacey, H. Kamal, G. Bogdanovic, S. Muschiol, D.J. Wullimann, T. Kammann, J. Emgård, Parrot, E. Folkesson, O. Rooyackers, L.I. Eriksson, J.I. Henter, A. Sönnerborg, T. Allander, J. Albert, M. Nielsen, J. Klingström, S. Gredmark-Russ, N.K. Björkström, J.K. Sandberg, D.A. Price, H.G. Ljunggren, S. Aleman, M. Buggert, and K.C.-S. Group. 2020. Robust T Cell Immunity in Convalescent Individuals with Asymptomatic or Mild COVID-19. Cell 183:158-168.e114.

Sercan, O., D. Stoycheva, G.J. Hämmerling, B. Arnold, and T. Schüler. 2010. IFN-gamma receptor signaling regulates memory CD8+ T cell differentiation. J Immunol 184:2855–2862.

SeyedAlinaghi, S., S. Oliaei, S. Kianzad, A.M. Afsahi, M. MohsseniPour, A. Barzegary, P. Mirzapour, F. Behnezhad, T. Noori, E. Mehraeen, O. Dadras, F. Voltarelli, and J.M. Sabatier. 2020. Reinfection risk of novel coronavirus (COVID-19): A systematic review of current evidence. World J Virol 9:79–90.

To, K.K., I.F. Hung, J.D. Ip, A.W. Chu, W.M. Chan, A.R. Tam, C.H. Fong, S. Yuan, H.W. Tsoi, A.C. Ng, L.L. Lee, P. Wan, E. Tso, W.K. To, D. Tsang, K.H. Chan, J.D. Huang, K.H. Kok, V.C. Cheng, and K.Y. Yuen. 2020. COVID-19 re-infection by a phylogenetically distinct SARS-coronavirus-2 strain confirmed by whole genome sequencing. Clin Infect Dis

Wang, D., B. Hu, C. Hu, F. Zhu, X. Liu, J. Zhang, B. Wang, H. Xiang, Z. Cheng, Y. Xiong, Y. Zhao, Y. Li, X. Wang, and Z. Peng. 2020. Clinical Characteristics of 138 Hospitalized Patients With 2019 Novel Coronavirus-Infected Pneumonia in Wuhan, China. JAMA 323:1061–1069.

Wu, Y., Z. Feng, P. Li, and Q. Yu. 2020. Relationship between ABO blood group distribution and clinical characteristics in patients with COVID-19. Clin Chim Acta 509:220–223.

Zhao, J., Y. Yang, H. Huang, D. Li, D. Gu, X. Lu, Z. Zhang, L. Liu, T. Liu, Y. Liu, Y. He, B. Sun, M. Wei, G. Yang, X. Wang, L. Zhang, X. Zhou, M. Xing, and P.G. Wang. 2020. Relationship between the ABO Blood Group and the COVID-19 Susceptibility. Clin Infect Dis

Zhou, C., Z. Huang, W. Tan, X. Li, W. Yin, Y. Xiao, Z. Tao, S. Geng, and Y. Hu. 2020. Predictive factors of severe coronavirus disease 2019 in previously healthy young adults: a single-center, retrospective study. Respir Res 21:157.

