## Supplemental Table and Figures for "ABO blood group is involved in the quality of the specific immune response"

### ABO group is associated with specific cellular and humoral immunity after COVID-19 infection

### Supplemental DATA

### Supplemental Table 1

| Whole blood labeling | | | |
|---|---|---|---|
| **Antigen** | **Fluorochrom** | **Company** | **City, Country** |
| CD3 | APC-Alexa Fluor700 | Beckman Coulter | Nyon, Switzerland |
| CD4 | VioBlue | Miltenyi Biotech | Bergisch Gladbach, Germany |
| CD8 | Viogreen | Miltenyi Biotech | Bergisch Gladbach, Germany |
| CD45 | PE-Cy5 | Becton Dickinson | Eysins, Switzerland |

| Specific celular T response assay | | | |
|---|---|---|---|
| **Antigen** | **Fluorochrom** | **Company** | **City, Country** |
| **Superficial labeling** | | | |
| CD4 | PE-Vio615 | Miltenyi Biotech | Bergisch Gladbach, Germany |
| CD8 | Viogreen | Miltenyi Biotech | Bergisch Gladbach, Germany |
| CD69 | R-PE-Cy5.1 | Beckman Coulter | Nyon, Switzerland |
| CD3 | APC/Cy5 | Beckman Coulter | Nyon, Switzerland |
| **Viability: Fixable viability Dye-eFluor450 (eBioscience, San Diego, CA, US)** | | | |
| **Intracellular labeling after cells were fixed/permeabilized with the Cytofix/Cytoperm solution kit (Beckton Dickinson, Eysins, Switzerland)** | | | |
| IL-2 | PE | Miltenyi Biotech | Bergisch Gladbach, Germany |
| TNFα | PE-Cy7 | eBioscience | San Diego, CA, US |
| IL-17A | FITC | eBioscience | San Diego, CA, US |
| IL-4 | APC | eBioscience | San Diego, CA, US |
| IFNγ | eFluor450 | eBioscience | San Diego, CA, US |

**Supplemental Table 1: antibodies used in flow cytometry analysis**

### Supplemental Figure 1

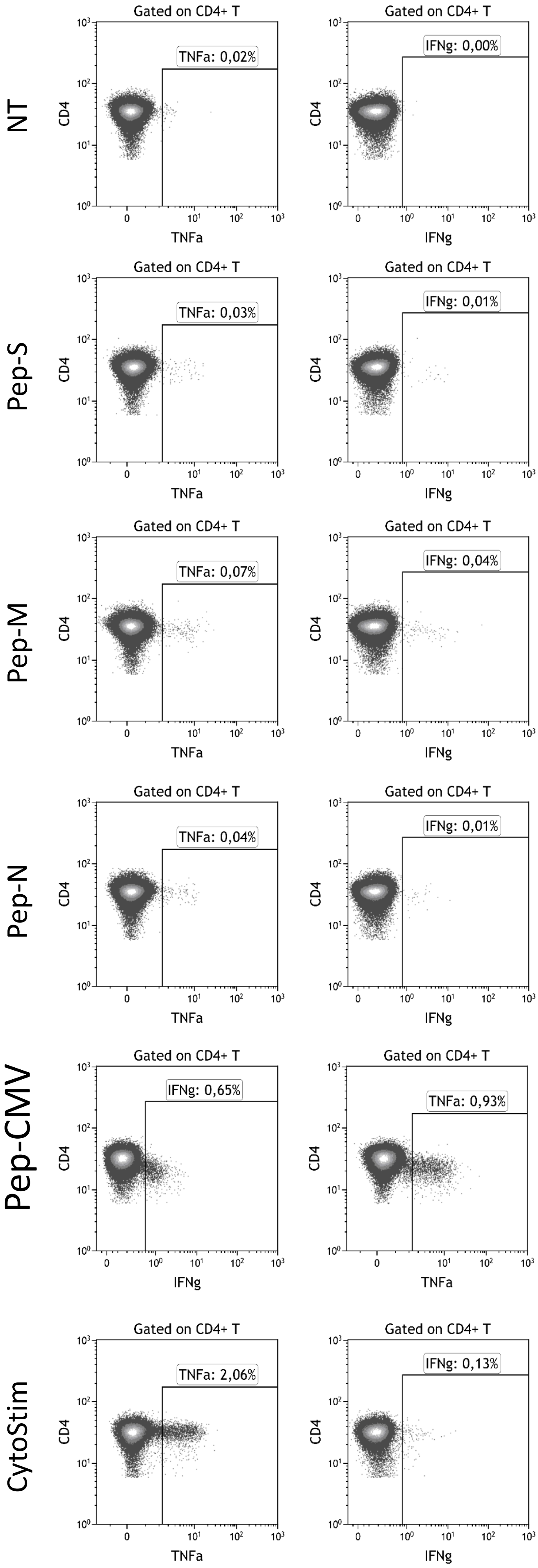


**Supplemental Figure 1: Specific CD4+ T-cell responses to SARS-CoV-2**

Representative examples of flow cytometry plots of specific IFN-γ- and TNF-α-producing CD4+ T cells for non-treated condition (NT) or after 6h of stimulation with Pep-S, Pep-M, Pep-N, Pep-CMV and CytoStim.

### Supplemental Figure 2

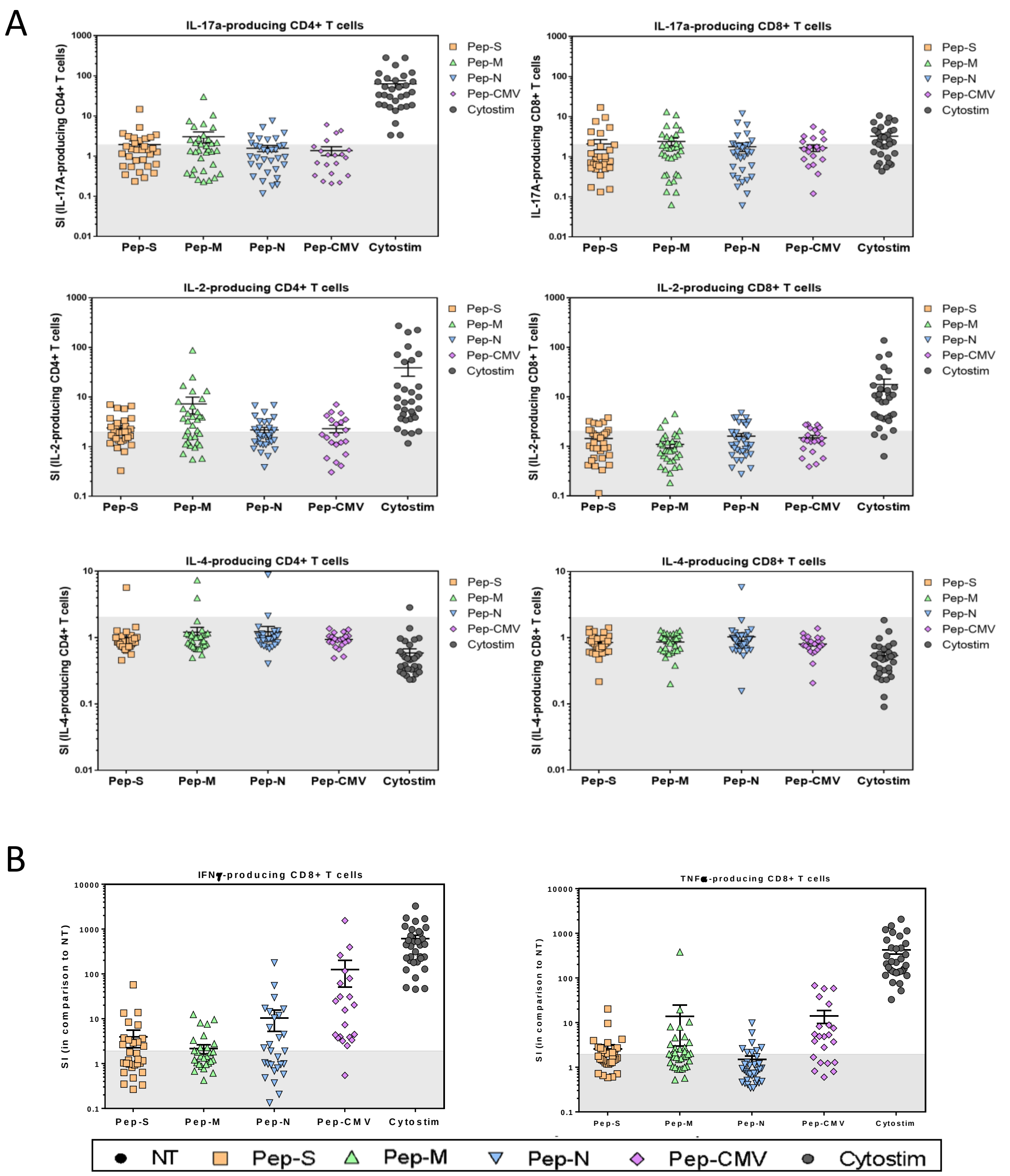


**Supplemental Figure 2: IL-17A, IL-2, IL-4, TNF-α and IFN-γ-specific T-cell responses to SARS-CoV-2-derived peptides pools**

PBMCs were isolated and stimulated by SARS-CoV-2-derived peptide pools (Pep-S; Pep-M and Pep-N), with CMV-derived peptides (Pep-CMV) or with CytoStim. (A) IL17-A, IL-2 and IL-4 expression was detected intracellularly by flow cytometry. Stimulation Index (SI) for each subjects was calculated dividing the frequency of cytoquine-producing CD4+ / CD8+ T cells in stimulated condition (Peptides and CytoStim) and frequency of cytokine-producing CD4+ / CD8+ T cells in the non-treated condition (NT). (B) SI of IFN-γ- and TNF-α-producing CD8+ T cells. Each symbol corresponds to an individual. The grey area represents the stimulation index inferior at 2 that is considered as a negative response to the stimulation in comparison to NT.

### Supplemental Figure 3

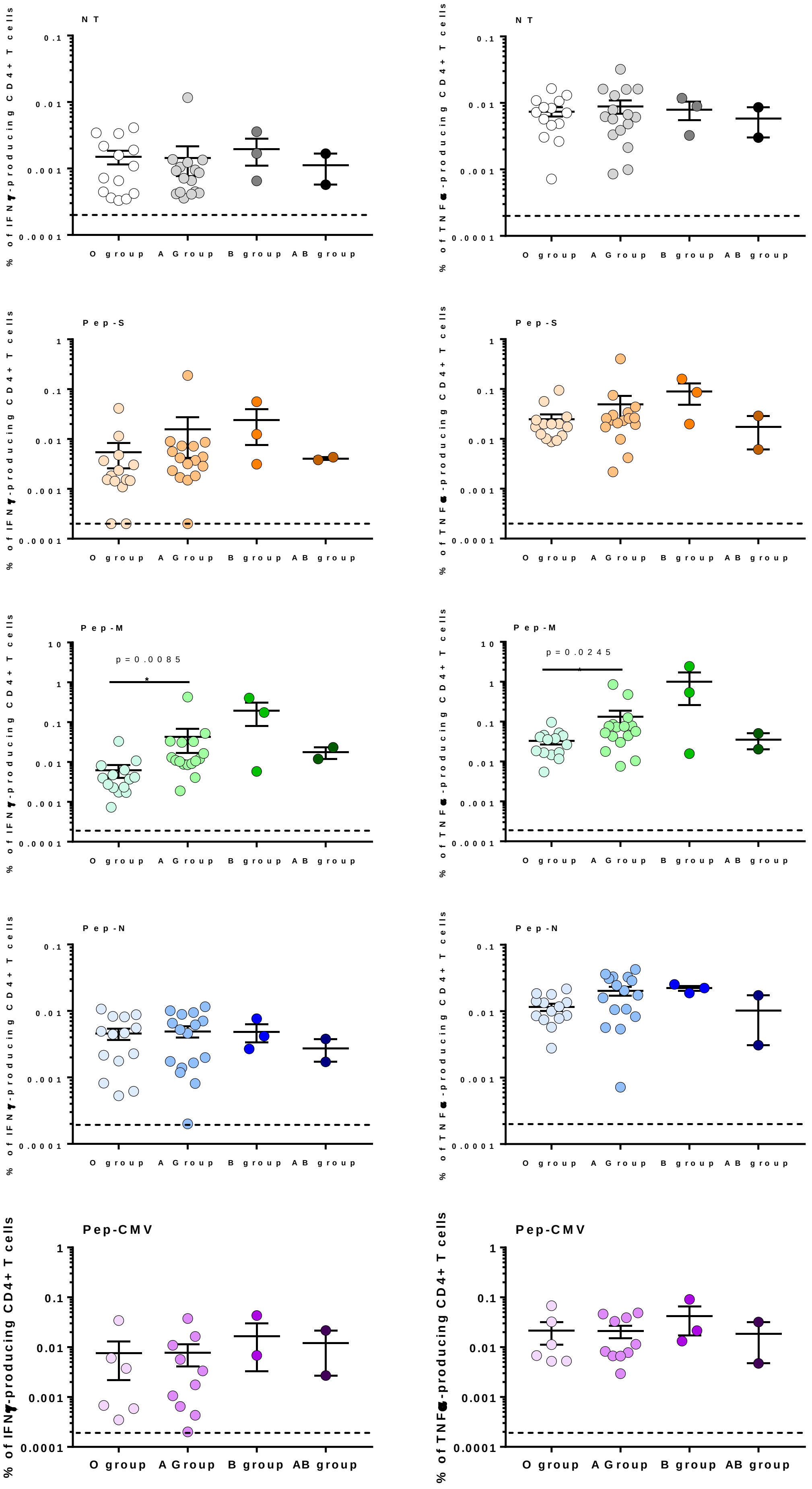


**Supplemental Figure 3: Blood groups as factor for CD4+ T cell specific TNF-α and IFN-γ responses**

Frequencies of IFN-γ- and TNF-α-producing CD4+ T cells in non-treated (NT) or stimulated PBMC with Pep-S; Pep-M; Pep-N and Pep-CMV in ABO groups. Symbols on the dotted line represent the frequency of specific CD4+ T cell with negative results. Mann-Whitney U tests. *p<0.05 was considered as significant.
